# Intracochlear pressure measurements quantify overall noise exposure to the inner ear during mastoidectomy

**DOI:** 10.1101/2025.03.30.646193

**Authors:** Emily J Bacalao, Nam K Lee, Juanantonio Ruiz, Brian Herrman, Nathaniel Greene

## Abstract

**Introduction:** The acoustic trauma patients are exposed to during otologic surgery is unclear, as previous studies have not captured the effects of vibrational noise on the inner ear. To fill this knowledge gap, we examined inner ear noise directly via intracochlear pressure changes during a mastoidectomy and cochleostomy.

**Methods:** Inner ear stimulation was measured using fiber optic pressure sensors inserted into the cochlea of cadaveric heads using a trans-canal approach. A tympanomeatal flap was elevated, cochleostomies were made into scala vestibuli and tympani, and sensors placed and secured to the cochlear promontory, and the flap resecured. Intracochlear pressures were measured with different drill bits, speeds, and during mastoidectomy and cochleostomy. The equivalent sound pressure level in the ear canal (L_Eq_) was calculated using the middle ear transfer function. Microphones were also placed in the surgeon’s ear, cadaver ear, and approximately three feet away.

**Results:** Results show sound pressure levels that could exceed 120 dBA (A-weighted dB SPL L_Eq_), with the highest levels observed during cochleostomy. Results showed broad band exposure, with a distinct peak at 1.33 kHz during the 80k RPM drill speed. Importantly, SPLs in the surgeon’s ear did not always predict patient noise exposure.

**Conclusion:** Surgically induced cochlear noise revealed levels high enough to potentially induce acoustic trauma, and sometimes at frequencies that may cause hearing loss not captured by a traditional audiogram (i.e. 1.33 kHz). This data characterizes surgical noise exposures, and will lead to permissible noise exposure guidelines and durations for both patients and surgeons.

## Introduction

Hearing loss has significant impacts on cognitive functioning, learning, communication, and quality of life (1), and accordingly, many otology and neurotology procedures are dedicated to improving or preserving a patient’s hearing. Mastoidectomy is one of the most common procedures done in otologic surgery as it provides access to the middle ear and inner ear structures through the mastoid bone. This procedure is used to treat diseases of the temporal bone, as well as access to the middle ear for cochlear implantation. Mastoidectomy requires removing the mastoid bone with a rotary surgical drill down to the level of the middle ear space (2) which can result in excessive noise exposure from drilling, suction, and blunt manipulation of the ossicles (3), but it remains unclear how much noise trauma drilling on the mastoid causes the patient.

Sensorineural hearing loss is generally considered rare, but the incidence and risks have been under investigation since at least the 1940’s (4). Hearing loss for mastoidectomy has been reported between 1.2-4.5% (5), and is often attributed to surgical mishaps or inflammatory processes. Reports of permanent threshold shifts after major ear surgery generally range from 0-5% (6-8), with a recent meta-analysis suggesting this risk is 3-5% (9), but some studies have reported substantially higher rates, with temporary threshold shifts of 48% at 2-4 kHz, and permanent shifts of 16% remaining 6 months after mastoidectomy (10). However, hearing loss rates during otologic procedures are thought to underestimate overall damage to the inner ear since standard audiometry cannot detect more subtle injuries. For example, extended high frequency hearing loss (>8 kHz) has been reported in 30% of ipsilateral ears, and 20% of contralateral ears following other skull base surgery (11). To increase understanding of hearing loss risk during otologic surgery, several studies have attempted to quantify patient noise exposures.

In contrast to a more traditional noise exposure, surgical drill exposure includes both air and bone conducted transmission to the inner ear. Air conducted transmission has been investigated by assessing the sound pressure level (SPL) adjacent to the patient’s ear, which reported only moderate exposures (84 dBA) (12). In contrast, probe-tube microphone measurements placed approximately 0.5 cm from the bone-drill interface or the round window in cadaver temporal bones varied between 116-131 dB SPL through different phases of mastoidectomy, including stapedotomy (13). These results suggest variable estimates dependent on microphone positioning.

To capture the vibratory bone-conducted noise, studies have used accelerometers on the mastoid or cochlear promontory. One study utilizing cadaver temporal bones estimated ipsilateral cochlear noise in excess of 100 dBA during drilling, and contralateral levels 5-10 dB lower (14), which varied with drill burr size, style, and drilling location (15). This bone conducted sound transmission has been reported to be maximal between 4-8 kHz for temporal bones (16), but has not been reported for whole heads, and vibratory measurements in patients undergoing surgery has been reported to be maximal between 2-4 kHz, approaching 117 dBA (17).

Importantly, no study has quantified the combined effect of both conduction pathways. These pathways interact via constructive and destructive interference, suggesting the overall exposure may be higher than either independently, resulting in more injury. To fill this gap, we measured intracochlear pressure (P_IC_) during surgical procedures, thereby measuring the combined exposure from both sound transmission pathways to the inner ear.

P_IC_ provide an ideal physical measurement of the sound transmitted to the inner ear since the difference in pressure across the basilar membrane correlates directly with cochlear microphonic in small animals (18), thus providing a direct measure of the driving force to the basilar membrane (19). Previous studies have quantified differential P_IC_ for normal and high level noise exposure (19,20), have quantified the effects of various inner ear pathologies (21), as well as blast exposure (22), which has both substantial air and bone conducted components. Importantly, P_IC_s have also been used to investigate the vibration of bony structures, such as the stapes or cochlear promontory and its effect on P_IC_ using pressure probes and 3D laser doppler vibrometer (23-26). Since the relationship between air and bone conduction pathways are not well understood, their interactions, particularly in the context of otologic surgery, have not been characterized to date.

Here, we utilize a novel trans-canal approach to directly measure P_IC_ induced by otologic surgery. *The goals of this study are to develop a technique for measuring P*_*IC*_*s during mastoidectomy surgery, and begin to characterize inner ear noise exposure of patients undergoing otologic surgery*. Our long term goals are to develop softer surgical techniques and drill designs, thereby lowering patient noise exposures, increasing surgical safety, and improving surgical outcomes.

## Materials and Methods

The trans-tympanic approach to the middle ear was developed and preliminary data collected from four hemicephalic cadaveric heads (four ears). The data presented here was collected from four fresh-frozen whole heads (eight ears) obtained from a medical donation program (MDG Global Inc.) with intact temporal bones and no history of middle ear disease. The use of cadaveric human tissue was in compliance with the University of Colorado Anschutz Medical Campus (CUAMC) Institutional Biosafety Committee (IBC #18-001), and the Colorado Multiple Institutional Review Board (COMIRB EXEMPT #18-2233).

### Temporal Bone Preparation

We have described the use of these pressure sensors several times previously (e.g. (3,25,27)). Here, we describe a novel surgical approach to implanting these electrodes via the external auditory canal (EAC).

Prior to each experiment, several checks were completed to ensure the specimen is representative of healthy normal living ears (e.g. (28)). Experiments were only conducted on specimens with no past medical history of middle ear disease or known anatomical anomalies, and external and middle ear anatomy was inspected prior to making measurements. Fresh-frozen cadaveric heads were defrosted and oriented to mimic intraoperative surgical positioning. As part of the inspection process the middle ear was accessed by raising the tympanic membrane via a tympanomeatal flap, carefully leaving annulus and tympanic membrane intact. The middle ear space was inspected, and the ossicular chain gently manipulated (to avoid high frequency transmission changes (29)) to ensure mobility and that a round window reflex was present, to ensure that no air bubbles were present in the cochlea. The round window niche and false membranes were removed to better expose the round window membrane.

The cochlear promontory was blue lined over scala vestibuli and scala tympani with a 0.5 mm diamond burr, and cochleostomies were opened, under water, with a fine straight pic. Fiber-optic pressure sensors (FOP-M260-ENCAP, FISO Inc., Quebec, QC, Canada) were routed into the middle ear cavity through the EAC, inserted into the cochleostomies, and sealed in place with alginate dental impression material (Jeltrate; Dentsply International Inc., York, PA) to record P_IC_. Fiber optic pressure sensors were secured to the cochlear promontory and bony canal wall, and the tympanomeatal flap was re-secured to the canal wall with cyanoacrylate adhesive. Pressure probe positioning on either side of the cochlear partition was verified after each experiment by dissecting the cochlear promontory bone between the two cochleostomies.

### Data Acquisition and Analysis

Data was collected via custom MATLAB (MathWorks Inc, Natick, MA, USA) software using a National Instruments (Austin, TX, USA) multifunction I/O data acquisition system (NIDAQ USB-6346). P_IC_s were measured with miniature fiber optic pressure sensors (FOP-M260-Encap; FISO Inc., Quebec, Canada) and digitized with an 8-channel fiber optic signal conditioner (Veloce-50; FISO). The sound pressure level (SPL) in the cadaver ear was measured with an Etymotic (Elk Grove Village, IL) ER-7C probe microphone, the SPLs in and at the entrance to the surgeon’s ear canal were measured with custom MEMS microphones (ARA Inc.; Littleton, CO, USA), and the SPL in the vicinity was monitored with a ¼” free-field microphone (G.R.A.S. 46B; Holte, Denmark). Notably, the patient ear canal was occluded by reflecting the pinna over the ear canal opening, providing a substantial attenuation for sound transmission to the tympanic membrane via the ear canal. Measurements were sampled at 96000 samples-per-second, and measurements made in 10s blocks.

P_IC_s were recorded beginning at the initiation of drilling, and concluded upon opening the inner ear via a cochleostomy. Recordings were scaled according to each sensor’s sensitivity, and a narrow-band notch filter applied at 60 Hz (and harmonic multiples of 4250 Hz for P_IC_) to remove artifacts. Middle ear transfer functions were estimated from prior estimates (20,29), and P_IC_ reverse filtered to estimate equivalent P_EAC_ (L_Eq_; Figure 2 top). An A-weighting filter was then applied to all channels (Figure 3 bottom) to facilitate noise exposure comparisons. Noise exposure measurements are reported as the root-mean-square (RMS) amplitude for each 10s recording, and aggregated across individual ears/experiments. Some recordings were additionally assessed by calculating the amplitude spectrum via the fast Fourier transform (FFT), and the time-frequency dependence assessed via a spectrogram. Statistical comparisons were conducted using the two-tailed, non-parametric Kruskal-Wallis and Mann– Whitney U tests, using α = 0.05. Follow-up post-hoc pairwise comparisons were completed using the Tukey-HSD test. All analyses were conducted using custom MATLAB software calling built-in functions.

**Figure 1.**
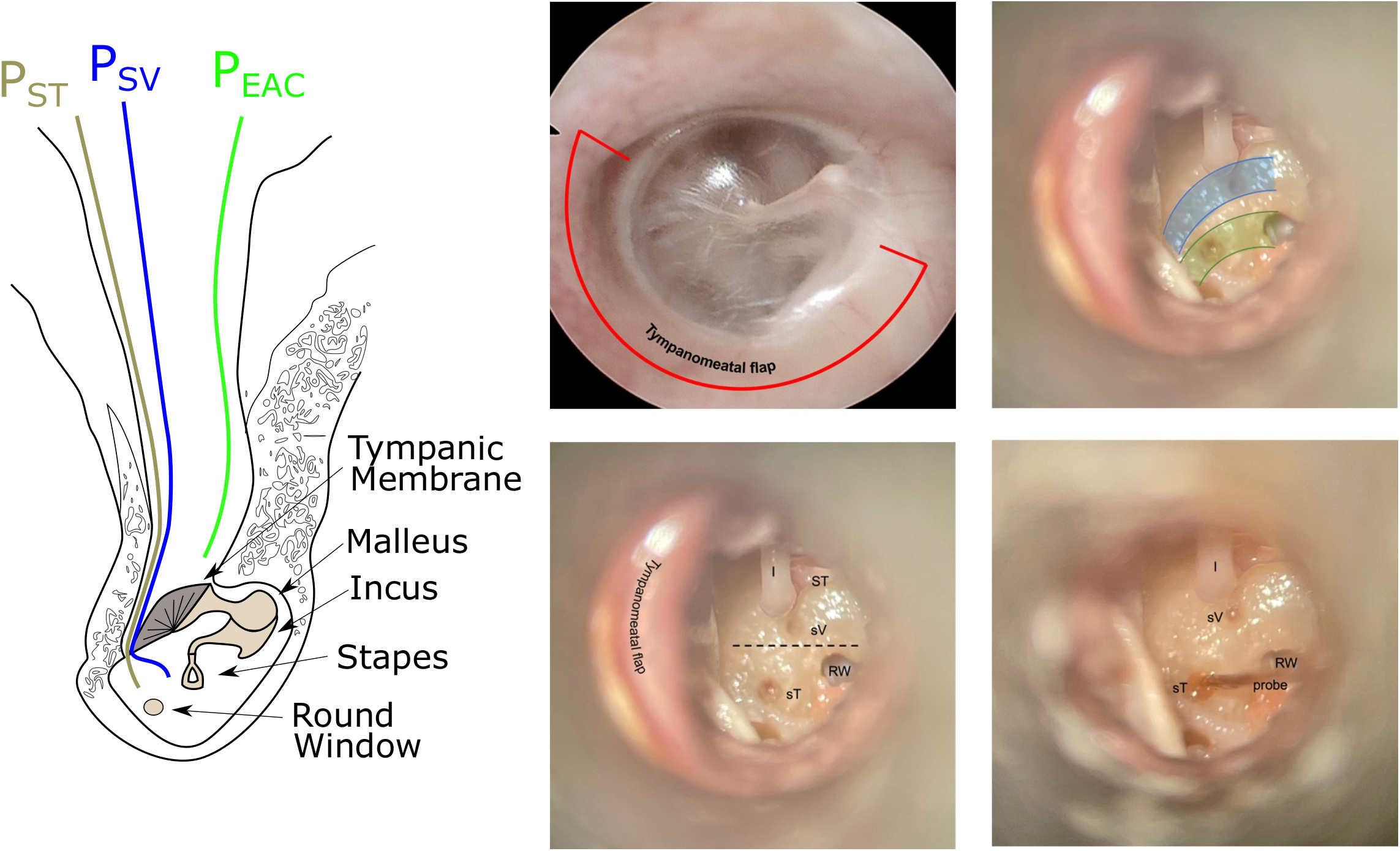
(color online). Left: An illustration representing the arrangement of a probe tube microphone (P_EAC_) into, and pressure probes passing through the patient EAC, under the tympanic membrane, and onto the cochlear promontory to access cochleostomies made into the scala vestibuli (P_SV_) and scala tympani (P_ST_). Right: (upper left) microscopic views of the intact tympanic membrane showing the location of the incision to make the Tympanomeatal flap; (lower left) the location of the Incus (I), stapes (ST), round window membrane (RW), and blue-lining over the scala vestibuli (sT) and scala tympani (sT), as well as the anticipated location of the cochlear partition (dashed line); (upper right) the anticipated course of the scala vestibuli (blue) and tympani (green); (lower right) the position of the scala tympani pressure probe after insertion.

**Figure 2.**
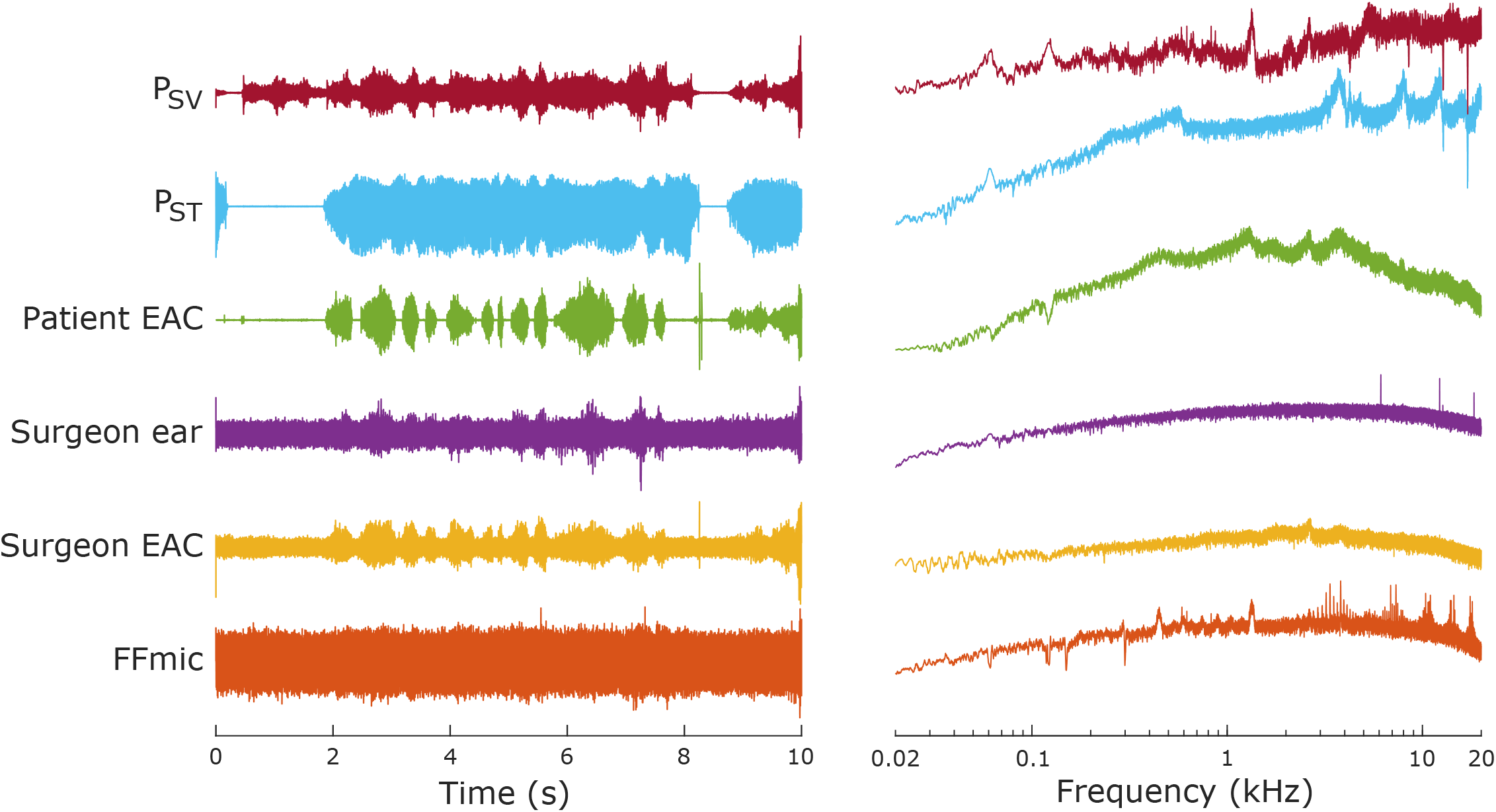
(color online). Example A-weighted recordings from each channel (left) and their corresponding amplitude spectra (right).

**Figure 3.**
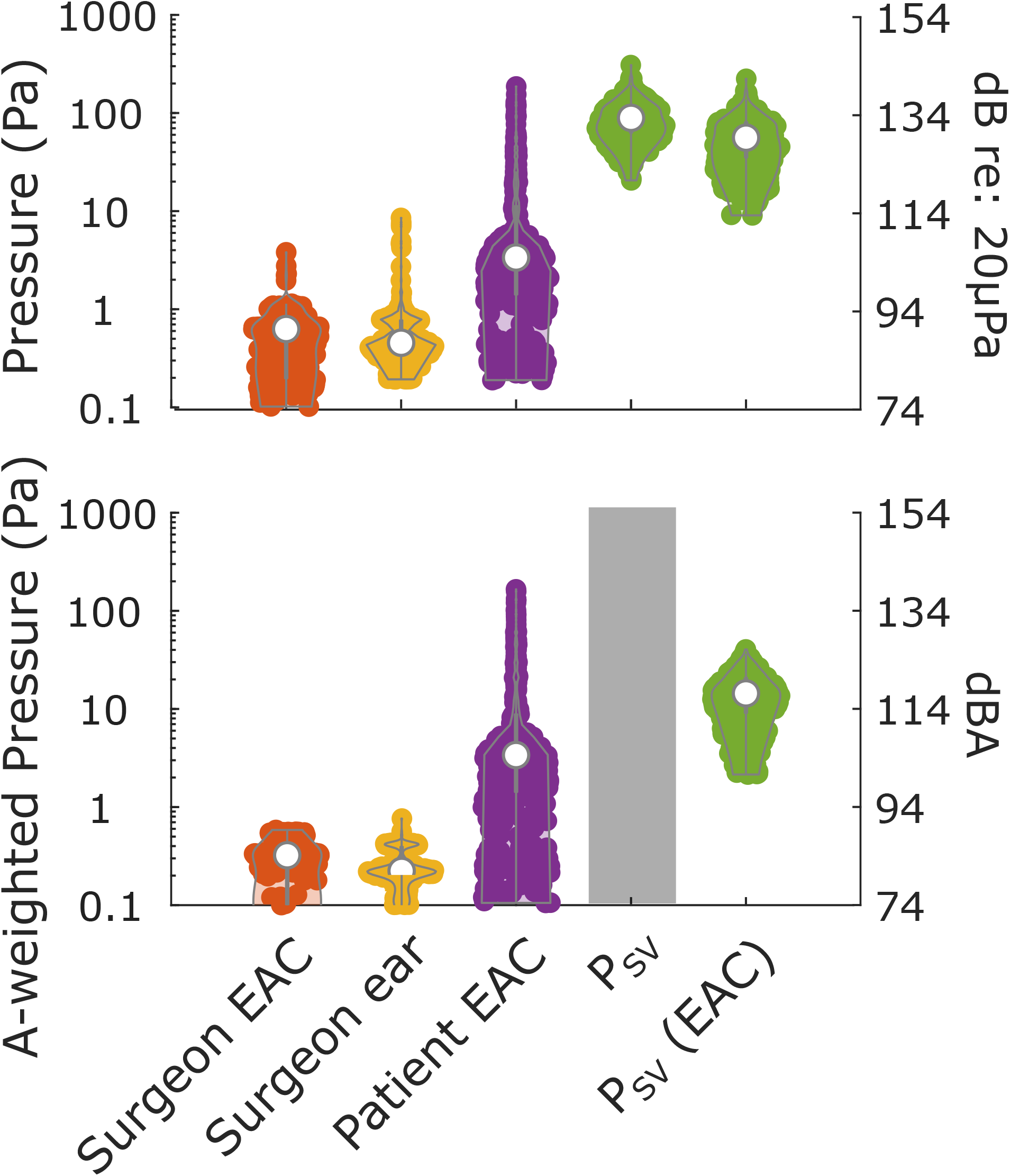
(color online). Top: violin plots of the root mean squared (RMS) amplitudes of recorded pressures from each sensor compared to scala vestibuli. Each filled point represents one 10s recording, and the colored outline displays the distribution of amplitudes recorded for each measurement. White circles show the median, and grey lines the interquartile range. Psv(EAC) represents the Psv as filtered through the middle ear transfer function to estimate the sound pressure level in the ear canal that would elicit the observe Psv (L_Eq_). Bottom: the same data after applying an A-weighting filter to estimate risk of hearing loss. Note, raw Psv data is excluded (gray bar) and only the estimated pressure in the ear canal (Psv(EAC)) is shown to facilitate comparisons to more typical air conducted sound exposures.

### Mastoidectomy

To simulate the mastoidectomy, a surgical incision is made 1 cm behind the auricle from the level of zygomatic root to the mastoid tip to ensure good exposure of the mastoid bone. The soft tissue is elevated and retracted with the same steps used intraoperatively. We then made measurements with a round 5mm cutting burr on mastoid cortical bone with drill speeds 20,000-80,000 rotations per minute (RPM) in intervals of 20,000 RPM. The mastoidectomy was begun using a round 5mm cutting burr at 80k RPM with a size 12-suction irrigator. Notations denoted when cortex, air cells, and antrum were encountered, and when facial recess and cochleostomies were opened. Mastoidectomy was generally performed with a drill speed of 80k RPM, as is typical for otologists at the University of Colorado Hospital, but the actual speed varied somewhat based on pedal input. Signals were monitored throughout the procedure.

When antrum was encountered the bur was switched to a 4mm coarse diamond bur to thin canal wall and define tegmen to better expose the incus in the fossa incudus. Once the canal wall was thinned the facial recess was opened with a 2mm coarse diamond bur and suction irrigation was downsized to size 5 suction irrigator. Upon entering middle ear through the facial recess, a 0.5mm diamond bit was used to make cochleostomies (smaller than standard to accommodate P_IC_ probes on the promontory). Once completed, the cochlea was opened to confirm probe placement in scala vestibuli and tympani.

## Results

Figure 2 shows example recordings for each sensor, including both P_IC_ probes (scala vestibuli: P_SV_ & scala tympani: P_ST_), microphone in the cadaver ear canal (Patient EAC), microphone in the surgeon’s ear canal (Surgeon EAC) and front of the ear canal opening (Surgeon ear), and a free field microphone (FFmic) adjacent to the recording hardware.

Significant electrical artifact was observed in the free field microphone throughout data acquisition thus are excluded from further analysis and consideration. P_IC_ changes in the scala tympani (P_ST_) and vestibuli (P_SV_) were noted throughout the simulated mastoidectomy.

Unfortunately the pressure probe in the scala tympani also showed high electrical artifact whenever the drill was activated, therefore those recordings are similarly excluded from further analysis, thus differential P_IC_ (P_Diff_) could not be calculated. However, clean responses free of artifacts were reliably recorded from the Surgeon ear & EAC, Patient EAC, and P_SV_.

Significant noise was generated whenever the drill contacted the skull, with distinct noise bursts appearing in all four channels simultaneously whenever the drill (which was running nearly continuously, see artifact in P_ST_ for timing) contacted the skull (Figure 2 Left). These noise bursts were present in all channels but most distinct in the Patient EAC, where external noise from external sources such as the suction was largely attenuated. Drill contact generated a persistent broadband noise exposure (Figure 2 Right), as well as a significant narrow band component at the frequency corresponding to the speed of the drill (80k RPM = 1.33 kHz).

To evaluate patient and surgeon exposures, the aggregate RMS amplitudes for Surgeon EAC and ear, Patient EAC, and P_SV_ are shown in figure 3. The top panel shows the unfiltered amplitudes from each measurement in addition to the equivalent ear canal pressure (L_Eq_) derived from P_SV_ (30). The bottom graph shows the same measurements with an A-weighting filter applied for comparisons to permissible noise exposure standards (e.g. NIOSH, OSHA).

Note, Psv is excluded to facilitate direct comparisons of airborne sound. Overall noise levels are reported in Table 1; most critically, A-weighted Psv(EAC) varied between the noise floor (∼100 dBA) and 126 dBA, with a median of ∼117, which was significantly different from each other recording (Chi-square=861, p<3*10^−186^, df=3); post-hoc pairwise comparisons reveal significant differences between each pair (p <<0.01), except Surgeon EAC and ear (p = 0.98).

**Table 1:** The minimum, median, and maximum RMS amplitude of A-weighted noise exposures recorded in each position.

| dBA | Minimum | Median | Maximum |
| --- | --- | --- | --- |
| Surgeon EAC | 67 | 84 | 89 |
| Surgeon Ear | 73 | 81 | 92 |
| Patient EAC | 74 | 104 | 138 |
| P<sub>SV</sub> | 100 | 117 | 126 |

Figure 4 compares A-weighted P_SV_ measurements based on drill type, drill speed, drill location and drill diameter. There are statistically significant differences between cutting (∼120 dBA) and diamond (∼114 dBA) burrs, but not across drill speeds (∼121 dB); however, comparisons are restricted due to limited data available. There was a difference based on location, with cortical bone drilling generating a median noise exposure of ∼120 dBA, which decreased to ∼114 dBA on the facial recess. There was similarly a strong difference across burr sizes, where exposure peaked at ∼120 dBA with the 4mm burr, and as only ∼112 dBA for the 2mm burr, though notably limited data was available for the 0.5mm burr.

**Figure 4.**
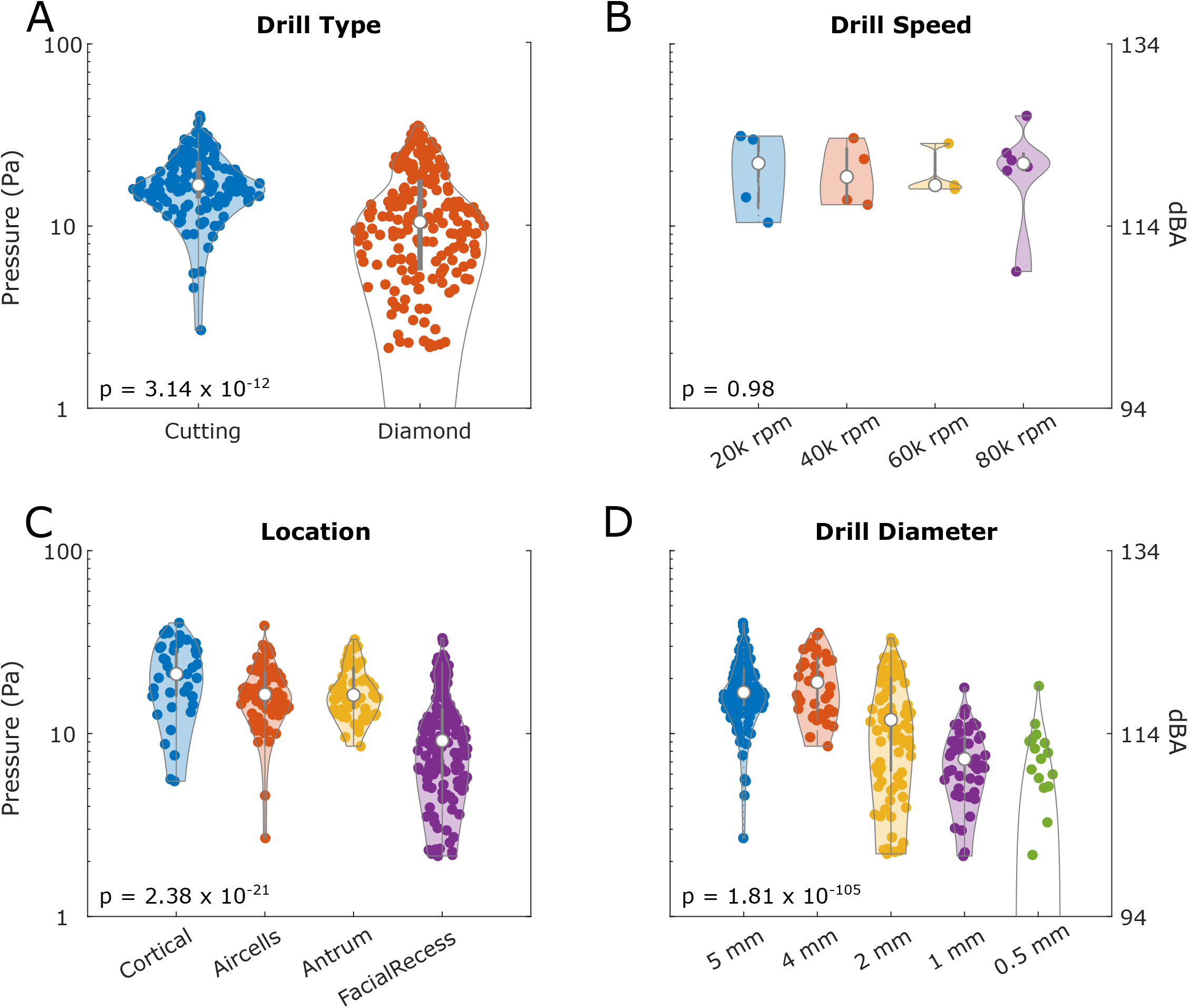
(color online). A-weighted measurement from P_SV_(EAC) sorted based on A) drill type, B) drill speed, C) drill location, and D) drill diameter. White circles represents each median and the grey bar represents the interquartile range. Each point represents results from a single 10s recording. P values are included representing the results from Mann-Whitney U (A), or Kruskal-Wallis (C-D) tests.

Figure 5 similarly compares A-weighted Surgeon ear measurements (just outside the ear canal entrance) based on drill type, drill speed, drill location and drill diameter. Compared to P_SV_ (figure 4), exposure levels are substantially lower and more variable; however, the trends observed across drill type, speed, location, and diameter are similar, with no significant differences across drill speeds, but significant differences across drill type, depth, and diameter.

**Figure 5.**
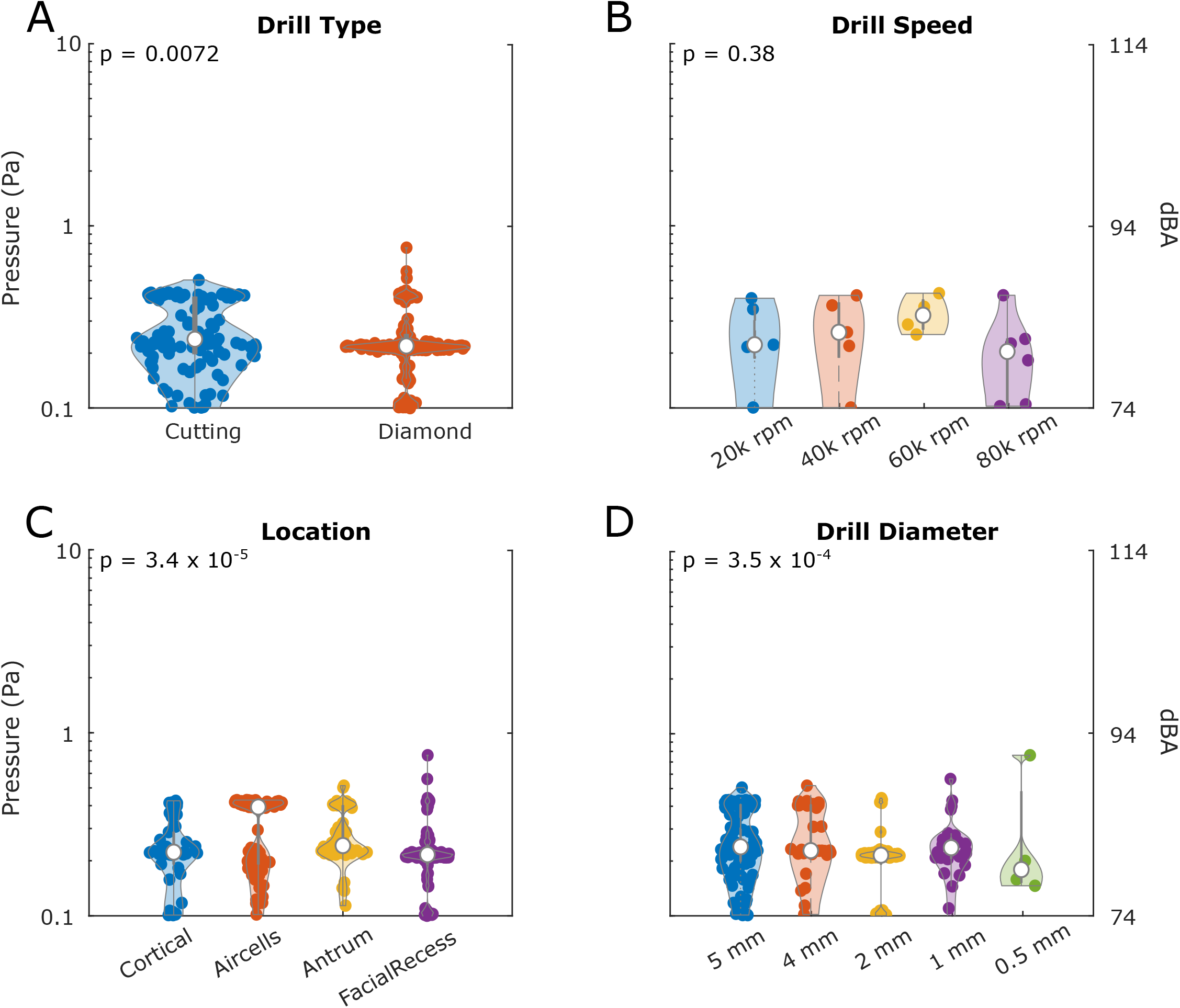
(color online). A-weighted measurement from Surgeon ear measurements sorted based on A) drill type, B) drill speed, C) drill location, and D) drill diameter. White circles represents each median and the grey bar represents the interquartile range. Each point represents results from a single 10s recording. P values are included representing the results from Mann-Whitney U (A), or Kruskal-Wallis (C-D) tests.

Since multiple factors covaried during measurements, the relative contributions of drill speed type, speed, location, and diameter, were further investigated using a multiple linear regression model fit to P_SV_ amplitudes. Results suggest a significant effect of drill diameter (tstat=8.0557, p<0.02*10^−14^) and type (tstat=3.572, p<5*10^−4^), but not location (tstat=1.429, p=0.154) or speed (tstat=-0.105, p=0.917). In contrast, a similar analysis on surgeon ear revealed significant effects of drill location (tstat=-3.8603, p<2*10^−4^) only, and not drill type (tstat=0.957, p=0.3395), diameter (tstat=-1.056, p=0.292), or speed (tstat=-1.496, p=0.136).These results suggest drill diameter and type drive patient inner ear noise exposures, and surgeon exposure does not necessarily predict patient exposure.

Figure 6 shows a spectrogram of the same recording shown in figure 2, in addition to time-domain (bottom) and frequency magnitude (right) graphs of this recording. Exposure induces a broad band noise exposure, revealed by vertical bands coincident with drill activity. Additionally, P_SV_ revealed a notable high pressure component at 1.333 kHz that corresponds with the frequency of the drill revolutions (80,000 RPM = 1,333 Hz) when the drill contacted the skull, revealing a notable spectral peak (right) not present when the drill pedal was not depressed (e.g. at t=0, 8.5s), or when the drill did not contact the skull (e.g. t=1s).

**Figure 6.**
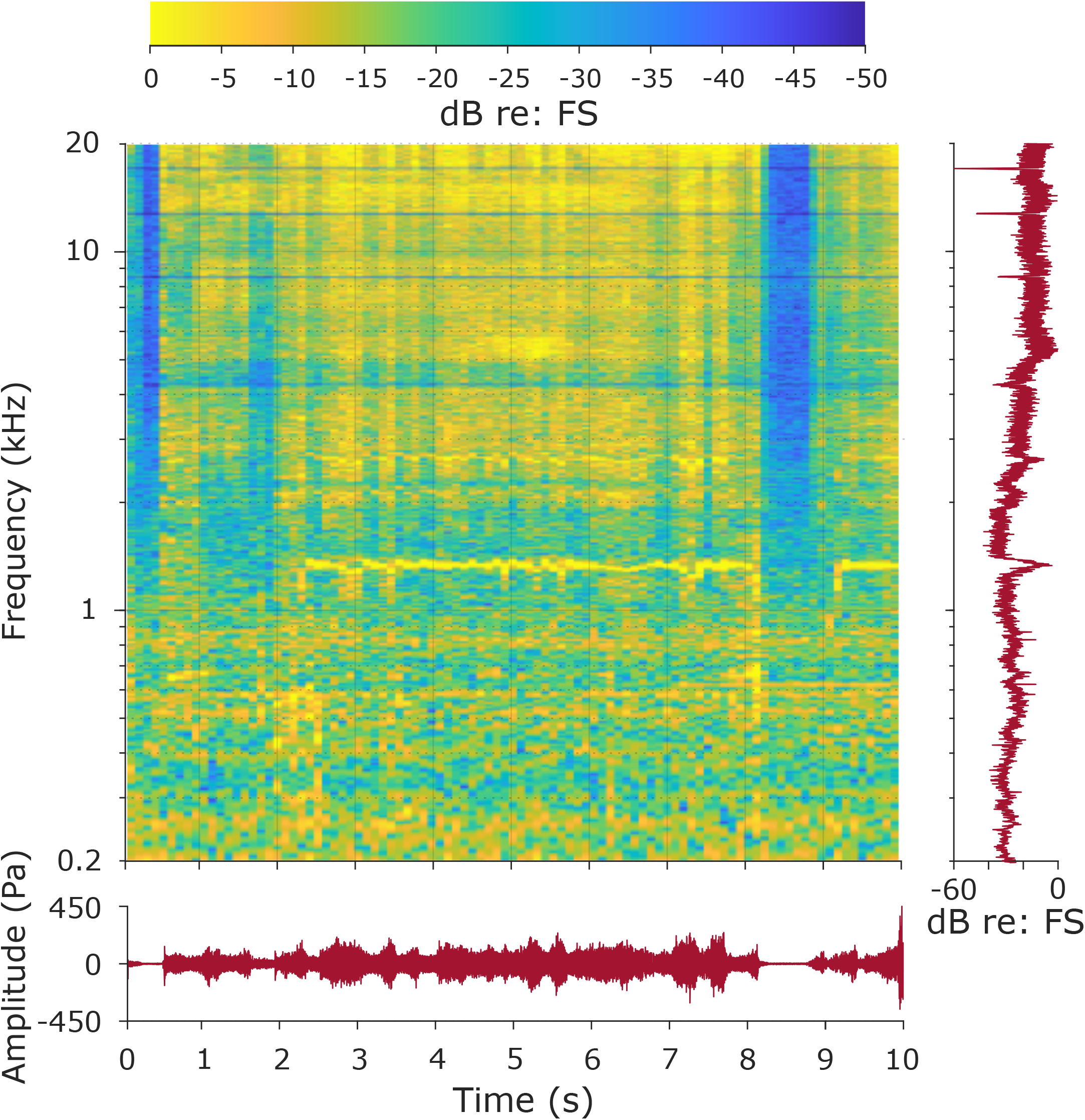
(color online). Spectrographic representation of the same representative P_SV_ recording during an 80k RPM drill application to the skull shown in Figure 2. Time is represented on the x-axis, amplitude spectrum on the y-axis, and sound power is represented by the color in each pixel (color bar at top, cooler colors represent lower amplitudes, in dB re: full scale). The time domain waveform is shown below, and the amplitude spectrum is shown to the right of the figure to illustrate the time and frequency dependence of the response amplitude.

The frequency content across multiple drill speeds are compared in Figure 7. A substantial component is observed at the drill speed (black arrow), as well as at several harmonics (gray arrows) at 80k RPM (bottom); however, these components are less prominent at lower drill speeds. A 1 kHz component is visible for 60k RPM, but no harmonics are visible, whereas no specific frequency component is visible at 40k RPM (666 Hz) or 20k RPM (333 Hz).

**Figure 7.**
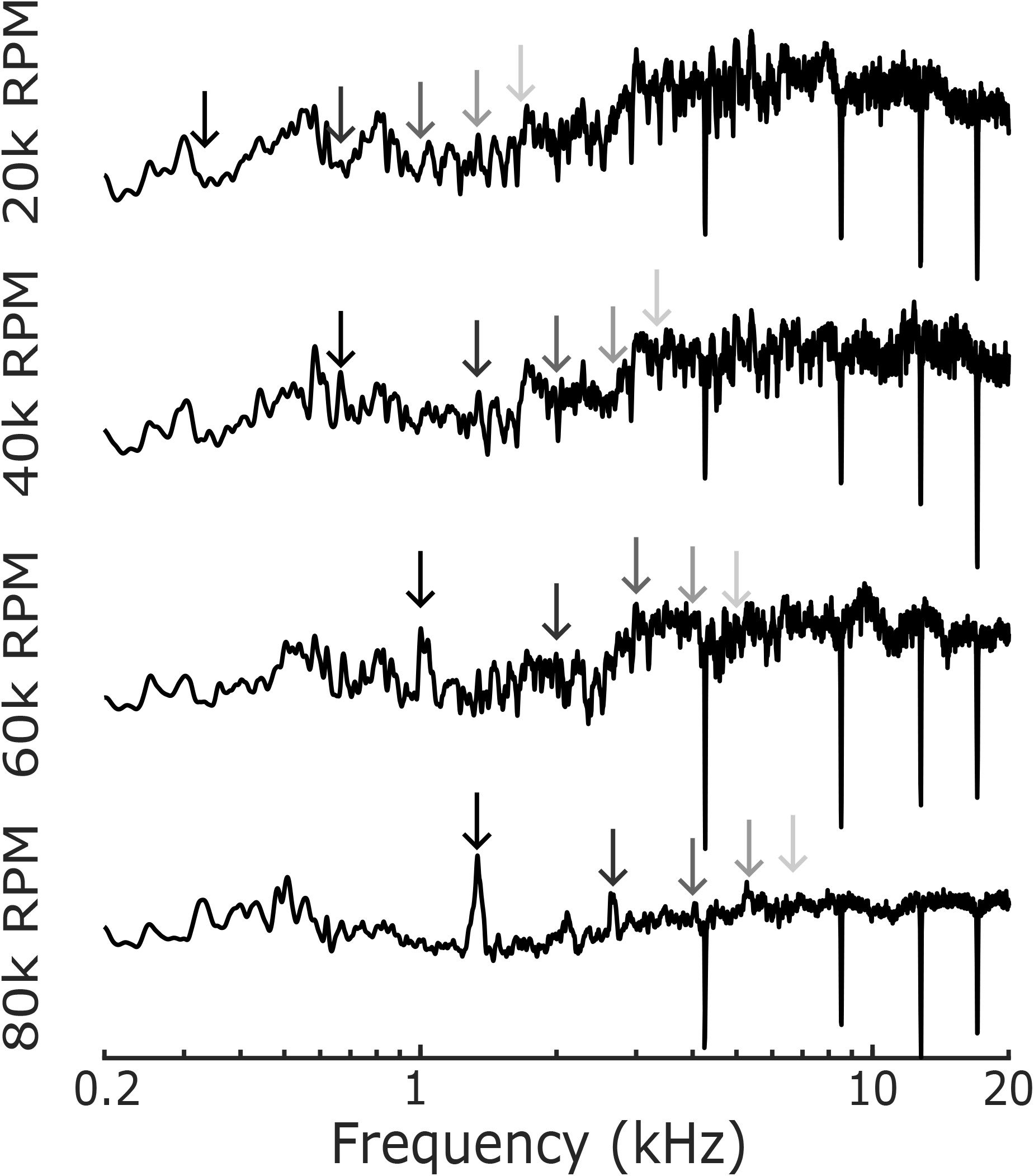
P_SV_ amplitude spectra from recordings during 20k, 40k, 60k, and 80k RPM drill applications to the skull. The frequency corresponding to the drill speed is represented by a black arrow, and the frequency of the first 4 harmonics are represented by grey shaded arrows in each. Note, the amplitude is reduced at ∼4, ∼8, ∼12, & ∼16 kHz as a result of notch filters removing electrical artifacts in each recording.

## Discussion

Results reveal substantial cochlear noise during mastoidectomy, with levels high enough to potentially induce acoustic trauma and hearing loss. Pressure levels in the cochlea were equivalent to noise exposures up to ∼126 dBA through various stages of the mastoidectomy.

These results are consistent with previous estimates reported in the literature, which can vary between 116-131 dB SPL (12-17). Additionally, these results reveal statistically significant differences in measurement locations, with median inner ear noise exposures approximately 13 dB higher than estimates from a microphone in the patient’s ear canal, suggesting that vibration substantially contributes to the overall inner ear noise exposure. Interestingly, ear canal noise exposures showed greater variability, with levels exceeding 130 dBA, suggesting that ear canal was a less reliable metric of overall noise exposure than P_IC_. Compared to occupational noise exposure limits from OSHA, which use a 5 dB exchange rate, permissible exposure limits are 130 dBA is less than 2 minutes per day (NIOSH limits, which use a 3 dB exchange rate are even shorter), suggesting lengthy surgical procedures place patients at considerable risk of inner ear trauma and hearing loss.

Furthermore, we observed significant differences in both surgeon and patient noise exposures based on drill type, location, and diameter. As predicted, cutting burrs were louder than diamond burrs, cortical locations were louder than facial recess, and large burrs were louder than small burrs, although the interdependence complicates the relationships between these factors. A multiple linear regression analysis was conducted to resolve this ambiguity, which suggested that drill type and diameter largely drove the patient’s noise exposure, whereas drill location predominantly dictated the surgeon’s noise exposure. Importantly, these results suggest that a surgeon’s perception is not a reliable predictor of patient noise exposure.

Our data additionally show a distinct narrow-band exposure at 1.333 kHz corresponding to the 80k RPM of drill speed. This narrow band noise exposure was not observed at lower drill speeds, which tended to show more broadband noise exposure, which may explain why previous studies utilizing slower drill speeds have not reported such a focal noise exposure.

Importantly, this frequency is not directly tested in a traditional audiogram, therefore its possible current rates of hearing loss stated in the literature after mastoidectomy may be an underestimation. Future studies should thus investigate hearing loss in post surgical patients at the specific frequencies induced by drill exposure, and not limit testing to the typical audiogram.

## Limitations

This study represents the first report utilizing a novel trans-tympanic approach to inserting pressure probes in the cochlea, thus several limitations must be considered when comparing results to prior studies. First, due to unforeseen electrical artifacts, we were unable to obtain reliable pressure recording from the operating room or scala tympani pressure sensors.

Since hearing perception is most directly correlated with the difference in pressure across the basilar membrane, and not the scala vestibuli pressure alone, we cannot provide a direct estimate of the acoustic energy delivered to the basilar membrane directly. Nevertheless, the P_SV_ measurements presented here represent the most direct measures of sound transmission to the inner ear available to date. Nevertheless, future studies should further refine this surgical and experimental approach to obtain P_Diff_ directly. Second, the trans-tympanic approach required removal, and replacement of the tympanic membrane, thus potentially altering the acoustics of the external and mechanics of the middle ear. Future studies should again further refine this technique to more closely approximate the unaltered condition, as well as confirm via acoustic and vibrometric measurements that the response of the tympanic membrane is consistent with an intact membrane, and unaltered by the probe insertion. Third, the results did not control for drill bit size, type, speed, or location. Recordings were made while a surgical resident conducted a mastoidectomy typical for the University of Colorado Hospital operating room. While the relative contributions of each are addressed via statistical comparisons, it is possible that a more systematic comparison across factors would yield different results.

Likewise, we did not consider additional drill factors, such as drill force, angle, irrigation, or surgeon experience, which may further complicate results. Finally, the cadaveric specimens were measured at room temperature, and not normal physiologic temperatures. Overall, despite the preliminary experimental approach and unfortunate technical difficulties, results from this study highlight the potential exposures, and risks, of surgically induced noise exposures, and highlights the need for further studies to improve surgical outcomes

## Conclusions

The current study reported inner ear noise exposure due to surgical drill contact with the skull for the first time, and suggests that the noise levels induced can be sufficient to induce hearing loss. Importantly, results suggest that the surgeon’s exposure is not a reliable indicator of the patient’s noise exposure, and that cumulative noise exposure should be considered when planning surgical procedures. Future work should further explore the impact of noise levels and durations on overall patient exposures during otologic procedures, and develop safer surgical techniques and contribute to safety guidelines.

## Acknowledgements

Dr. S Cass and S Gubbels for helpful discussions and development of the surgical approach, Dr. O Kalmanson for videos demonstrating mastoidectomy.

## Notes

### Competing Interest Statement

The authors have declared no competing interest.

## References

1. Kuo CY, Huang BR, Chen HC et al. Surgical results of retrograde mastoidectomy with primary reconstruction of the ear canal and mastoid cavity. Biomed Res Int 2015;2015:517035.

2. Flint PW, Francis HW, Haughey BH et al. Cummings otolaryngology : head and neck surgery. Philadelphia, PA: Elsevier, 2021:3 volumes.

3. Banakis Hartl RM, Mattingly JK, Greene NT et al. Drill-induced Cochlear Injury During Otologic Surgery: Intracochlear Pressure Evidence of Acoustic Trauma. Otol Neurotol 2017;38:938–47.

4. Rüedi L, Furrer W. Das akustische Trauma. Practica Oto-Rhino-Laryngologica 1946;8:504-.

5. Tos M, Lau T, Plate S. Sensorineural hearing loss following chronic ear surgery. Ann Otol Rhinol Laryngol 1984;93:403–9.

6. Tos M, Lau T, Plate S. Sensorineural hearing loss following chronic ear surgery. Annals of Otology, Rhinology & Laryngology 1984;93:403–9.

7. Urquhart A, McIntosh W, Bodenstein N. Drill-generated sensorineural hearing loss following mastoid surgery. The Laryngoscope 1992;102:689–92.

8. Palva T, Kärjä J, Palva A. High-tone sensorineural losses following chronic ear surgery. Archives of Otolaryngology 1973;98:176–8.

9. Yücel L, Satar B, Serdar MA. Meta-analysis of hearing outcomes of chronic otitis media surgery in the only hearing ear. Auris Nasus Larynx 2022;49:322–34.

10. Kazikdas KC, Onal K, Yildirim N. Sensorineural hearing loss after ossicular manipulation and drill. Ear, Nose & Throat Journal 2015;94:9.

11. Thirumala P, Meigh K, Dasyam N et al. The incidence of high-frequency hearing loss after microvascular decompression for trigeminal neuralgia, glossopharyngeal neuralgia, or geniculate neuralgia. Journal of Neurosurgery 2015;123:1500–6.

12. Man A, Winerman I. Does drill noise during mastoid surgery affect the contralateral ear? Am J Otol 1985;6:334–5.

13. Yin X, Stromberg AK, Duan M. Evaluation of the noise generated by otological electrical drills and suction during cadaver surgery. Acta Otolaryngol 2011;131:1132–5.

14. Kylen P, Arlinger S. Drill-generated noise levels in ear surgery. Acta Otolaryngol 1976;82:402–9.

15. Kylen P, Stjernvall JE, Arlinger S. Variables affecting the drill-generated noise levels in ear surgery. Acta Otolaryngol 1977;84:252–9.

16. Hilmi OJ, McKee RH, Abel EW et al. Do high-speed drills generate high-frequency noise in mastoid surgery? Otol Neurotol 2012;33:2–5.

17. Spencer M, Reid A. Drill-generated noise levels in mastoid surgery. The Journal of Laryngology & Otology 1985;99:967–72.

18. Dancer A, Franke R. Intracochlear sound pressure measurements in guinea pigs. Hear Res 1980;2:191–205.

19. Nakajima HH, Dong W, Olson ES et al. Differential intracochlear sound pressure measurements in normal human temporal bones. J Assoc Res Otolaryngol 2009;10:23–36.

20. Greene NT, Jenkins HA, Tollin DJ et al. Stapes displacement and intracochlear pressure in response to very high level, low frequency sounds. Hear Res 2017;348:16–30.

21. Pisano DV, Niesten ME, Merchant SN et al. The effect of superior semicircular canal dehiscence on intracochlear sound pressures. Audiol Neurootol 2012;17:338–48.

22. Greene NT, Alhussaini MA, Easter JR et al. Intracochlear pressure measurements during acoustic shock wave exposure. Hear Res 2018;365:149–64.

23. Borgers C, Fierens G, Putzeys T et al. Reducing Artifacts in Intracochlear Pressure Measurements to Study Sound Transmission by Bone Conduction Stimulation in Humans. Otol Neurotol 2019;40:e858–e67.

24. Stieger C, Guan X, Farahmand RB et al. Intracochlear Sound Pressure Measurements in Normal Human Temporal Bones During Bone Conduction Stimulation. J Assoc Res Otolaryngol 2018;19:523–39.

25. Mattingly JK, Banakis Hartl RM, Jenkins HA et al. A Comparison of Intracochlear Pressures During Ipsilateral and Contralateral Stimulation With a Bone Conduction Implant. Ear Hear 2020;41:312–22.

26. Dobrev I, Pfiffner F, Roosli C. Intracochlear pressure and temporal bone motion interaction under bone conduction stimulation. Hear Res 2023;435:108818.

27. Misch ES, Banakis Hartl RM, Gubbels SP et al. Risks of Intracochlear Pressures From Laser Stapedotomy. Otol Neurotol 2020;41:308–17.

28. Koch M, Essinger TM, Maier H et al. Methods and reference data for middle ear transfer functions. Sci Rep 2022;12:17241.

29. Nakajima H, McHugh C, Ravicz M et al. A mystery solved: human high-frequency middle-ear motion. Res Sq 2024.

30. Rosowski JJ, Chien W, Ravicz ME et al. Testing a method for quantifying the output of implantable middle ear hearing devices. Audiol Neurootol 2007;12:265–76.

